# Systematic Review on the Role of Microbial Activities on Nutrient Cycling and Transformation Implication for Soil Fertility and Crop Productivity

**DOI:** 10.1101/2024.09.02.610905

**Authors:** Tesfaye Bayu

**Affiliations:** Department of natural resources management Debre Markos university Burie campus, Burie, Ethiopia

**Keywords:** Crop productivity, Nutrient cycling, Nutrient transformation, Soil fertility

## Abstract

Soil microorganisms play a vital role in the regulation of the transformation and cycle of soil nutrients, thereby improving soil fertility and crop productivity. These microbes, associated with plants, contribute significantly to plant growth and development by improving nutrient cycling and crop productivity by improving soil fertility. This systematic review aims to assess the impact of microbial activity on nutrient cycle and transformation, which includes soil fertility and crop productivity improvement. The PRISMA flow methodology systematically included articles from various geographic regions. Through analyzing 120 articles, this review sought to address the question at hand. Among the articles analyzed, 31.4% indicate that soil microbial activity directly regulates nutrient cycling, while 68.6% suggest that microbial activity enhances soil fertility and crop productivity. The systematic review concludes that microbial activity has a significant effect on nutrient cycle and transformation, as well as on improving soil fertility and crop productivity. Farmers, policymakers, and experts are encouraged to manage soil microorganisms to regulate nutrient cycling, directly influencing soil fertility and crop productivity, thus promoting sustainable agricultural development.

## 1. Introduction

Microbial communities are crucial in shaping soil structure (Delmont et al., 2011), and play a vital role in supporting various ecosystem services in terrestrial environments, such as enhancing plant productivity, purifying water, and storing carbon (Torsvik and vres, 2002). Soil, especially the rhizosphere surrounding plant roots, hosts a diverse array of microorganisms (Chamkhi et al., 2022), many of which form symbiotic relationships with plants, thus creating integrated systems (Mustafa et al., 2019). Despite their small size, soil microbes significantly impact biogeochemical cycles involving essential nutrients such as carbon, nitrogen, sulfur, and phosphorus, as well as other mineral cycles (Griffiths, 1994). Furthermore, they contribute to soil health by producing antibacterial compounds that suppress soil pathogens, thereby enhancing plant disease resistance (Haas and Défago, 2005).

The nutrient cycle operates as an interconnected system, encompassing sulfur, phosphorus, nitrogen, carbon, and other essential elements that cycle simultaneously (Azcón-Aguilar and Barea, 2015). Soil organic matter is critical to provide carbon-based nutrients to microorganisms and maintain humus levels (King, 2011). Humus is essential for soil structure and acts as a reservoir for slowly available nutrients such as nitrogen, phosphorus, and sulfur, facilitated by microbial activity (Blackwell, 2021). Soil microbes play a significant role in the extraction of nitrogen, sulfur, and phosphorus from organic matter in agricultural soils (Walter, 1959). Ammonification is conducted predominantly by specific prokaryotes, including bacteria and archaea, using the enzyme nitrogenase (Olivares, Bedmar, and Sanjuán, 2013). Recent studies suggest that nitrogenase-like sequences in microbial genomes are more widespread than previously recognized (Santos et al., 2012). Additionally, soil bacteria and fungi can degrade organic phosphorus compounds by mineralization, releasing orthophosphate into the soil solution (Richardson et al., 2009).

Soil microorganisms play a vital role in the cycling of nutrients by activating their genes and extracellular enzymes (Borchard et al., 2019). They enhance various processes such as mineralization, solubilization, oxidation, reduction, nitrification, denitrification, and fixation, thus increasing soil nitrogen availability and aiding plant nitrogen uptake (Zhang et al., 2021). Specific microbes facilitate the release of less accessible phosphorus forms through organic acid exudation (for example, malic, oxalic and citric acids) and phosphatase enzymes, making phosphorus more usable (Richardson et al., 2009; Yu et al., 2021).

Soil microbes play a crucial role in supporting plant growth and development, improving nutrient cycling, and increasing crop productivity (Li et al., 2017). The composition of soil microbial populations significantly influences soil fertility and its ability to support crop growth (Shah et al., 2021). Interactions between plants and microbes are pivotal in ecosystem functioning, with the dynamics influenced by nutrient availability (Jiao et al., 2021). Microbes are extensively used to stimulate plant growth by regulating plant processes and aiding in nitrogen fixation (Ahmad et al., 2008).

Microbial-based biofertilizers are essential in agriculture, improving crop productivity and promoting sustainable agroecosystems (Ahirwar et al., 2019). These products consist of various microbial-based formulations crucial for enhancing biological processes in the plant-microbe-soil system (Singh et al., 2016). Bacteria and fungi with plant growth-promoting traits play a key role in the production of effective biofertilizers (Smith et al., 2011). Beneficial rhizosphere microorganisms contribute to plant growth by influencing biochemical pathways, such as regulating plant hormones, suppressing pathogens, and improving soil nutrient availability (Jacoby et al., 2017).

Direct mechanisms involve improving resource acquisition (eg nitrogen, phosphorus, potassium, micronutrients), modulating plant hormones, and producing molecules such as siderophores and aminocyclopropane-1-carboxylate deaminase, which reduce ethylene levels and alleviate osmotic stress in plants (Nadeem et al., 2014). Indirect mechanisms include the biocontrol of phytopathogens through antimicrobial metabolites, nutrient competition, and inducing systemic resistance (Munees and Mulugeta, 2014; Planchamp et al., 2015). These interactions can improve the resilience of plants to pathogens. This review explores the role of soil microorganisms in nutrient cycling, transformation, soil fertility, and crop production (Planchamp et al., 2015).

Exploring how bacterial and fungal inoculants interact with organic fertilizers is promising for improving soil fertility and structure (Hooda et al., 2023). This strategy could lead to the development of environmentally friendly, chemical-free soil nutrient management systems that meet global food demands and rejuvenate degraded soils. Using both bacteria and fungi alongside organic amendments offers a refined approach to sustainable soil fertility management and enhanced crop production (Sagar et al., 2021), especially in severely degraded soils.

This systematic review aims to investigate the relationship between microbial activity and nutrient cycle and transformation, highlighting its effects on soil fertility and productivity. It also proposes effective soil management strategies. The decline in soil fertility is increasingly evident due to globalization, population growth, environmental degradation, and climate change. The primary objective of this paper is to evaluate how microbial activity impacts nutrient cycle and transformation, both essential to maintain soil fertility and to increase plant productivity. The review has two specific goals: to assess the contribution of microbial activity to nutrient cycling and to analyze its role in nutrient transformation. This review is significant because managing soil fertility and improving crop productivity are critical issues, especially in the context of climate change. By suggesting sustainable soil management strategies, it aims to address these challenges effectively. Understanding the complex relationship between microbial activity and nutrient cycling can offer valuable insights to policymakers and stakeholders aiming to improve soil fertility and crop yields.

The review used the PRISMA flow methodology to systematically include articles from diverse geographic regions. The PRISMA model follows six crucial steps for selecting relevant and valuable articles. This method was chosen to prioritize recent and pertinent research on climate change and invasive species, enhancing the effectiveness and quality of the review. By adhering to the PRISMA model, the review demonstrates a clear and transparent approach to its composition.

This comprehensive review explores the relationship between microbial activity and nutrient cycle and transformation, assessing their effects on soil fertility and crop productivity. The review is organized into four main sections. The first section investigates the impact of microbial activity on the cycling of nutrients, and the second section examines its role in the transformation of nutrients. The third section discusses the interaction between microbial activity and soil fertility in different regions. The final section evaluates the global landscape of microbial activity and its potential implications for future crop productivity trends.

## 2. Methods

### 2.1. Search strategy, keywords and selection criteria

A systematic review of the literature was conducted following the PRISMA framework developed by Koutsos, Menexes, and Dordas (2019). The review process involved six steps: scoping, planning, identification and search, selection of publications, eligibility evaluation, and presentation and interpretation. These steps were meticulously followed to maintain rigor and transparency. The review adhered to the PRISMA statement protocol (Moher et al., 2009), recognized as the gold standard for systematic reviews, to ensure scientific rigor and a transparent evaluation.

The reviewer selected appropriate databases and developed search strategies to include only peer-reviewed, high-quality articles from reputable sources. The search was limited to specific databases, including WOS, SCOPUS, Google Scholar, and publications from global organizations. A search string with relevant keywords and Boolean operators was created to address the themes of climate change and invasive species. The search string, “TI = (microbial activity* AND nutrient cycling* OR microbial activity and nutrient transformation* AND microbial activity * OR * AND soil fertility*) AND TS = (microbial activity)” was used to refine the search results.

In March 2023, the investigation began, leading to the selection of 1,126 articles for inclusion in the study. These articles underwent a thorough selection process to ensure they met predetermined criteria, including publication in peer-reviewed scientific journals, English language usage, and classification as “original articles.” Furthermore, selected publications should focus on soil microbial activity, nutrient cycling, and transformation, with an emphasis on soil fertility and crop productivity.

From the initial selection of 1,126 articles, 726 were excluded because they focused on different aspects, such as the effects of soil fauna and flora, including nematodes and trees, or soil fertility management for agricultural development. Full-text articles that did not meet the predetermined criteria were also excluded, leaving a total of 232 articles. After a thorough analysis, 120 articles were ultimately included in the review. These articles were classified into four groups: 1 Nutrient cycling and soil microbial activity.2 Nutrient transformation and soil microbial activity, 3 Soil fertility and soil microorganism activity, and 4 Crop productivity and microbial activity. An article was assigned to one of these categories if it primarily addressed the respective theme.

The selected articles were analyzed, summarizing details such as publication year, geographic scope, methodology, approach, and reference data used. Due to variations in objectives, scientific backgrounds and data sources among the articles, minor nuances in theoretical and methodological approaches were not extensively discussed, in line with the nature of this review. The reviewer then established logical connections between various study fields and sustainable development goals (SDGs) based on the recent literature and their understanding of the subject. To aid in this process, the collected articles were classified by research field. It is important to note that the summaries and supporting citations are not directly related to the articles included in the systematic review of the literature. Figure 1 illustrates the framework used in this review.

**Figure 1.**
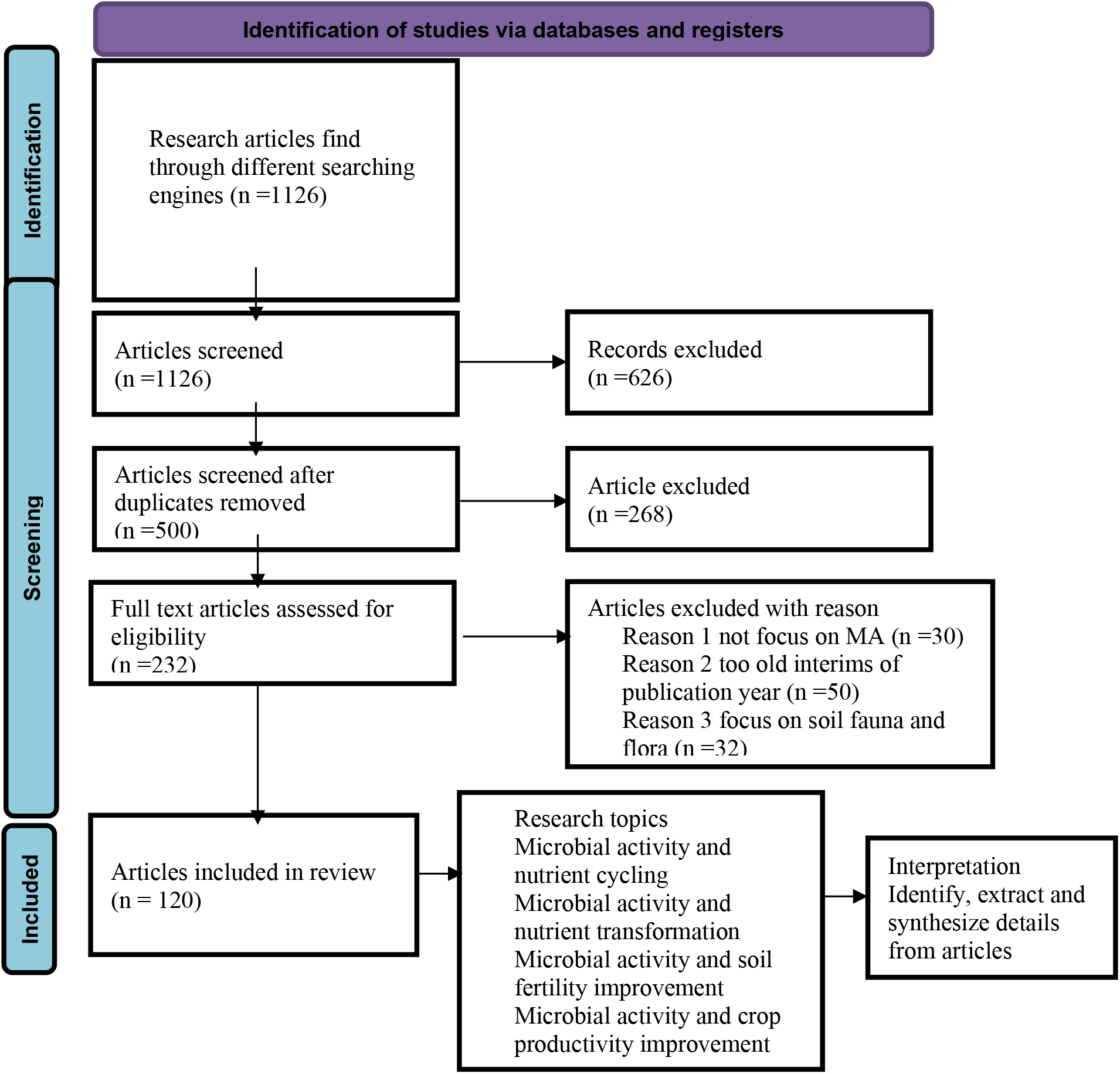
Systematic review framework based on the PRISMA principle

## 3. Results

### 3.1. Methodological, thematic, temporal and spatial distribution of literature

The research collected data from the literature on the effects of soil microbial activity on nutrient cycling, transformation, soil fertility, and crop productivity. The literature was organized and analyzed to understand the global impact of soil microbial activity on these processes. Texts were classified into four categories according to their content: 1 How microbial activity enhances nutrient cycling, 2. How soil microorganisms mediate nutrient transformation, 3. How soil microbial activity improves soil fertility and 4. How microbial activity increases crop productivity (see Figure 1).

Several reviewed articles highlight the involvement of microbial activity in the regulation of nutrient cycling and transformation (see Figure 3). These studies emphasize that soil microbial activity significantly impacts soil fertility and crop productivity worldwide. Quantitative data analysis methods were predominant in more than half of the reviewed studies (51.4%), while mixed methods were used in 27% and qualitative methods in 21.6% (see Figure 2). This diversity in data analysis approaches underscores the review’s goal of encompassing publications using various techniques to enhance overall quality.

**Figure 2.**
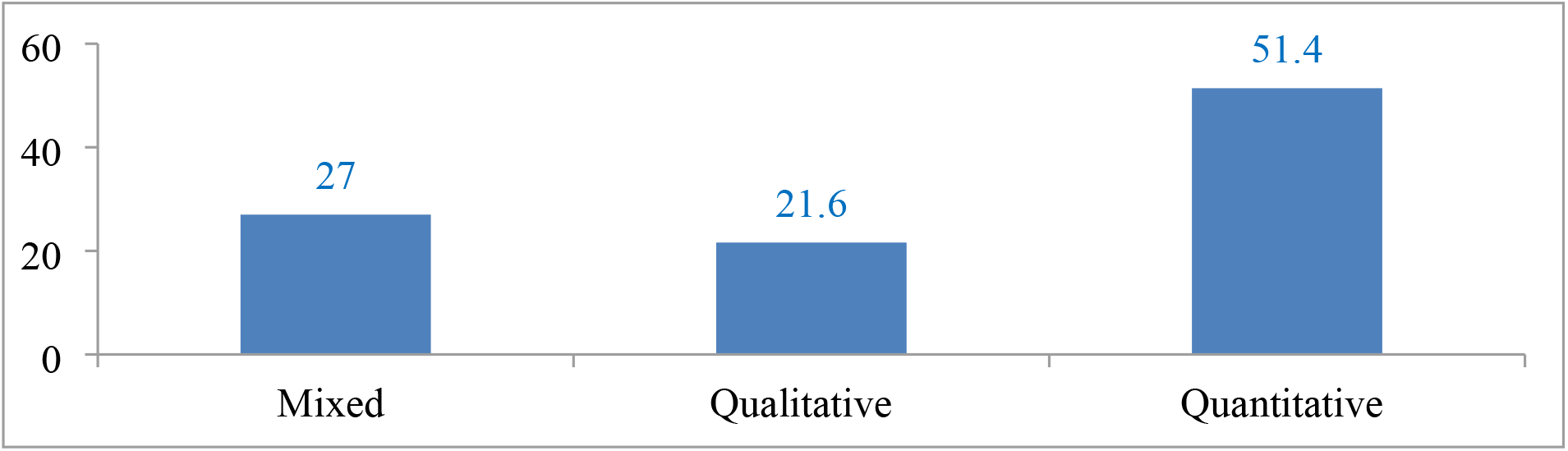
Distribution of articles according to the model used for the analysis

**Figure 3.**
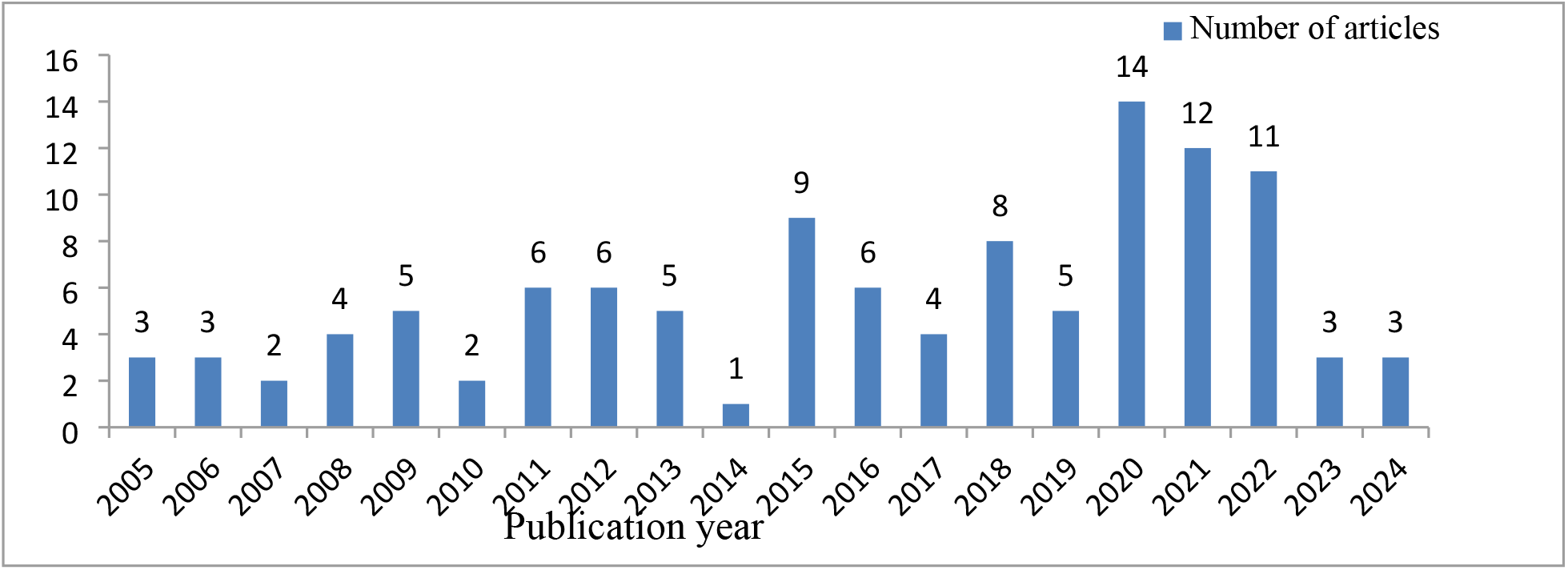
Temporal distribution of articles

The analysis indicates that six articles were evenly distributed in 2011, 2012, and 2016. The year 2020 had the highest number of publications (14), followed by 12 in 2021 (see Figure 4). Furthermore, Figure 4 illustrates five publications each in 2019, 2013, and 209. The fluctuating trend depicted in the chart highlights the limited growth in published articles in this field. This underscores the importance of ongoing research and publication on the role of soil microbial activities in nutrient cycling, transformation, soil fertility enhancement, and crop productivity in various regions.

**Figure 4.**
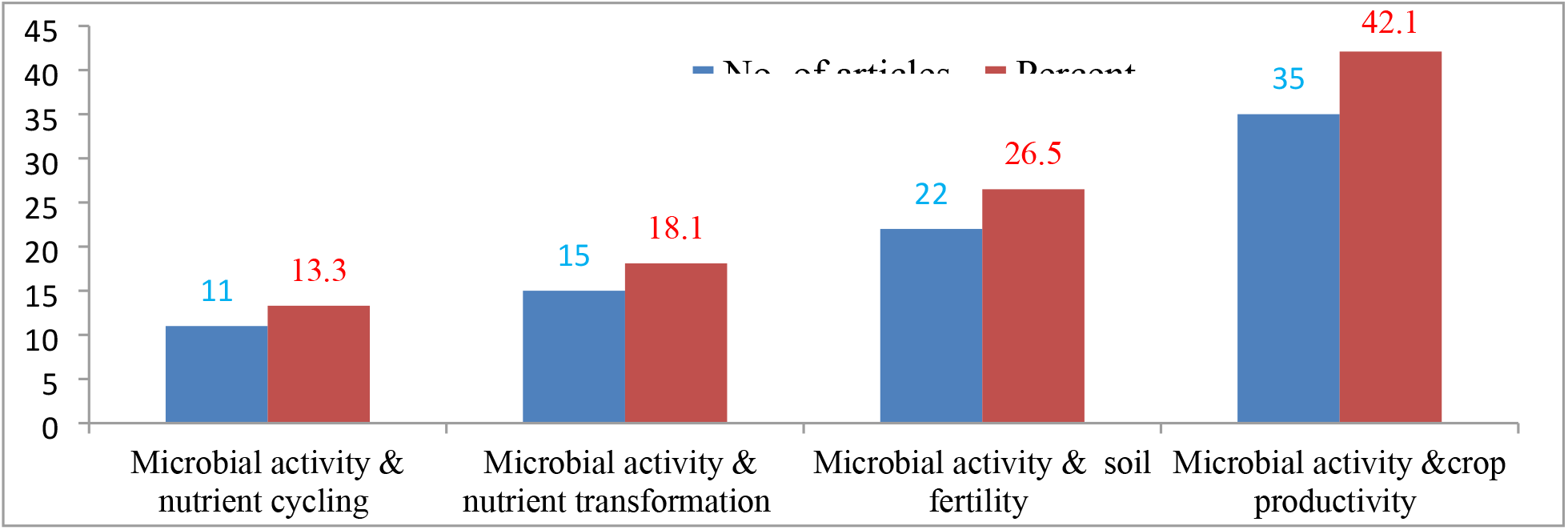
Number of articles per searching keywords

The 120 selected articles fall into four main research domains, as depicted in Figure 1. During the selection process, the research on soil microbial activity and its impact on crop productivity was the most prevalent, accounting for 42.1% of the articles reviewed. Those addressing soil microbial activity and soil fertility improvement made up 26.5%, while articles focusing on microbial activity in relation to nutrient cycle and transformation represented 18.1% and 13.3%, respectively (see Figure 5).

**Figure 5.**
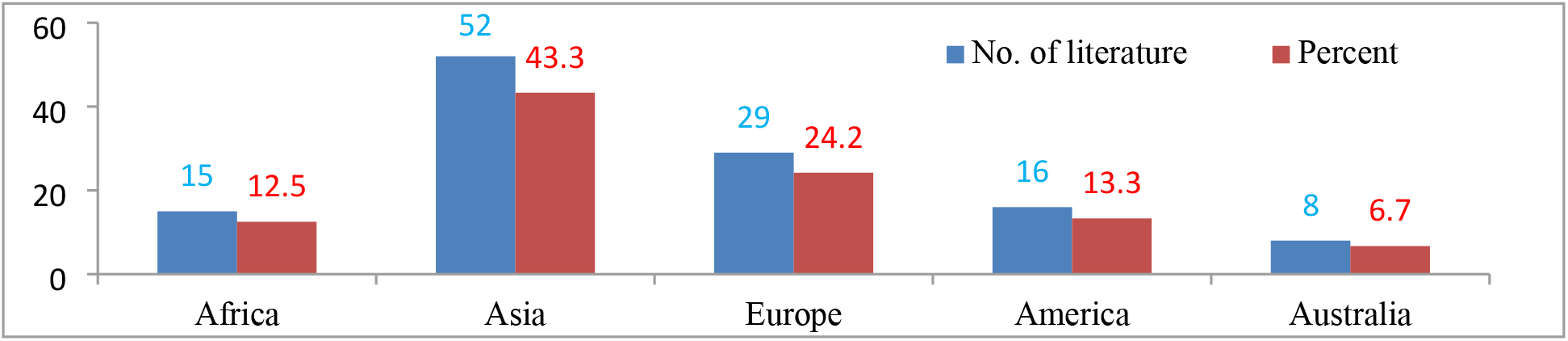
Geographic distribution of articles

The study is divided into four sections that examine the interaction between microbial activities and nutrient cycling and transformation, and its implications for improving soil fertility and crop productivity (see Figure 6). The review includes articles from five distinct regions, with the majority originating from Asia (43.3%) and Europe (24.2%) (Figure 6). A smaller portion of the literature comes from Australia (6.7%) and Africa (12.5%).

## 4. Discussion

### 4.1 Role of microbial activities in nutrient cycling

A small proportion of the articles (13.3%) indicate that microbial activity affects the cycling of nutrients (see Figure 4). For example, Javahery and Rokhzadi (2011) found that efficient strains of Azotobacter, Azospirillum, Phosphobacter, and Rhizobacter play a significant role in improving nitrogen availability for nitrogen cycling. Similarly, Walter (1959) demonstrated that soil microorganisms are vital for returning carbon dioxide to the atmosphere through decomposition and respiration, both on land and in the sea, which is essential for plant growth.

Microbial solubilization of mineral phosphorus (P) and the mineralization of organic P by microorganisms are both crucial in the cycling of P. Organic P, mainly in the form of inositol polyphosphates, can constitute 30-50% of total soil P (Li et al., 2015). This mineralization is driven by P-hydrolyzing enzymes such as phytases and phosphatases, produced primarily by fungi and bacteria (Huang et al., 2021). Acid phosphatases, which dephosphorylate phosphorus compounds and phosphoanhydride bonds in organic compounds, are key in mineralization of P (Alori, Glick and Babalola, 2017). Beyond enhancing P bioavailability, these microorganisms produce phytohormones, increase resilience to biotic and abiotic stress through the synthesis of compounds such as antifungal agents, and regulate vital metabolic pathways (Sharma et al., 2013).

In addition to phosphate-solubilizing microorganisms (PSM), arbuscular mycorrhizal fungi (AMF) also play a vital role in phosphorus cycling within various agrosystems. AMF form the most widespread symbiotic relationships on Earth, involving over 80% of terrestrial vascular plants. This symbiosis involves bidirectional nutrient exchange (Wang, Liu, and Zhu, 2018). Their role in the mobilization and uptake of phosphorus can be substantial, contributing up to 80% of the total uptake of phosphorus, depending on the conditions of the soil and the treatments with phosphorus (Adhya et al., 2015). Schnepf and Roose (2005) used a mathematical model to quantitatively assess the contribution of AMF hyphae to phosphorus acquisition by plants, demonstrating that plants may rely entirely on the mycorrhizal pathway for phosphorus nutrition, thus influencing the soil phosphorus cycle.

Morphologically, AMF extends the root system through its mycelium network, allowing plants to access nutrients beyond the rhizosphere. This mechanism is especially significant in phosphorus-deficient conditions, improving phosphorus cycling (Wang et al., 2017). AMF provides an additional phosphorus uptake pathway, with the arbuscular interface acting as the symbiotic interface. This process bypasses direct uptake by the root epidermis, which often becomes obstructed due to the rapid formation of a depleted zone from rapid phosphorus absorption from the soil solution (Shen et al., 2011).

Plants serve as the main carbon source for soil microorganisms by providing carbon-rich root exudates produced during photosynthesis and through input of plant residues (Kaiser et al., 2015). In return, soil microorganisms facilitate the cycling of various soil nutrients, including carbon, nitrogen, phosphorus, calcium, and magnesium, by fixing environmental elements and decomposing organic matter. For example, Grzyb, Wolna-Maruwka, and Niewiadomska (2021) highlight that microorganisms are essential in the nitrogen transformation cycle due to their genes encoding enzymes involved in nitrogen metabolism, which indicate the potential of the cycle.

### 4.2 Role of microbial activities in the transformation of nutrients

Figure 4 from a review study shows that 18.1% of the studies analyzed report a correlation between soil microorganism activity and soil nutrient transformation. For example, Murphy et al. (2007) found that the oxidation of H2S, the most reduced sulfur compound in nature, to S and potentially to SO42-, is performed by three types of microorganisms: Beggiatoa, Thiobacillus, and photoautotrophic bacteria from Rhodobacteriineae. The first two types obtain both the energy and the reduction power (hydrogen supply) needed for the assimilation of CO2 from the oxidation reaction, while Thiorhodaceae (purple and green sulfur bacteria) gain energy from photoautotrophic reactions, using reduced sulfur compounds as the hydrogen donor (Murphy et al., 2007). Additionally, Beura et al. (2022) observed that microorganisms contribute to phosphorus transformation by solubilizing mineral phosphates through various mechanisms: (i) generating carbon dioxide and organic acids, (ii) reducing ferric phosphates to more soluble ferrous compounds, and (iii) producing hydrogen sulfide, which enhances the solubility of phosphates and is a specific case of (ii). Soil fauna and microorganisms play crucial roles in the conversion of dead organic material to simple organic compounds and minerals. For example, Satyanarayana et al. (2012) highlighted the involvement of microorganisms such as Pseudomonas, Bacillus, Micrococcus, *Flavobacterium, Fusarium, Sclerotium, Aspergillus*, and *Penicillium* in the solubilization process, a critical step in phosphorus transformation. A strain of phosphate-solubilizing bacteria, NII-0909 from Micrococcus sp., exhibits polyvalent properties, including phosphate solubilization and siderophore production, making phosphorus available for plant uptake (Dastager, Deepa, and Pandey, 2010). Hongyuan et al. (2015) isolated *Aspergillus fumigatus* and *A. niger* from decaying cassava peels, demonstrating their ability to transform cassava waste into phosphate biofertilizers through semisolid fermentation. *Burkholderia vietnamiensis*, known for its tolerance to stress, produces gluconic and 2-keto gluconic acids, contributing to the transformation of phosphate (Nautiyal et al., 2000). *Enterobacter* and *Burkholderia*, isolated from sunflower rhizospheres, produce siderophores and indolic compounds (IC), which aid in the solubilization and transformation of phosphate into a form readily available for plant uptake and cycling (Beneduzi, Ambrosini and Passaglia, 2012). Additionally, potassium-solubilizing microorganisms (KSMs) such as genera Aspergillus, Bacillus and Clostridium are effective in solubilizing and transforming potassium in the soil, thus improving its mobilization in various crops (Bahadur et al., 2016).

Microorganisms primarily sustain themselves by transforming carbon through oxidation and reduction processes. For example, aerobic methane-oxidizing bacteria facilitate carbon transformation in oxygen-rich environments, such as the soil surface and aerobic layers of wetlands (Nercessian et al., 2005; Ma et al., 2020). On the contrary, in waterlogged, oxygen-deprived soils, CO_2_ reduction occurs through hydrogenotrophic archaea and methanogenic bacteria (Lu and Conrad, 2005). Soil microorganisms also play a crucial role in the formation of soil organic matter (SOM), a significant reservoir of soil organic carbon (SOC). The decomposition processes driven by these microorganisms involve various lytic enzymes, such as amylase, glucosidase, proteases, cellulase, chitinase, and phenol oxidase that break down complex macromolecules into simpler compounds. These compounds are then either assimilated by microbial communities or further transformed into CO_2_ to generate energy (Burns et al., 2013).

Bacteria play a vital role in enzymatic transformations within soil ecosystems. For example, Azospirillum, a microaerobic bacterium, supports nitrogen fixation in association with grass roots. Nitrification, the conversion of ammonia to nitrite ions, is conducted by nitrifying bacteria such as Nitrosomonas, while Nitrobacter oxidizes nitrites to nitrates. Clostridium pasteurianum, an obligate anaerobe, converts atmospheric nitrogen into ammonia, enriching soil with nitrogen. Research has highlighted that the presence of red alder (Alnus rubra) improves nitrification processes primarily mediated by prokaryotes, underscoring autotrophic nitrifiers as the main sink for NH4 + in soils (Sharma et al., 2023).

Moreover, bacterial fermentations produce acidic by-products that convert insoluble phosphates into soluble forms, thereby promoting plant growth. Vascular arbuscular mycorrhizae, working alongside plant roots, also contribute to the conversion of insoluble phosphates into soluble forms. Specific bacteria such as Thiobacillus ferrooxidans and iron bacteria of the Gallionella genus have the ability to oxidize ferrous (Fe^2+^) iron into ferric (Fe^3+^) iron (Anderson et al., 2006).

### 4.3 Role of microbial activities in improving soil fertility

According to the reviewed literature (see Figure 4), 26.5% of the studies indicate that microbial activity significantly impacts soil fertility. For example, research has shown that soil microorganisms can exploit, translocate, mineralize, and mobilize reserves of phosphorus (P), potassium (K) and iron (Fe), as well as improve organic matter content and fix nitrogen from the atmosphere (Kafle et al., 2019). In addition, the structure and function of microbial communities in soils profoundly influence soil matrices, including chemical and physical properties such as the quality and quantity of organic matter of soil, the pH, and the redox conditions. Merino-Martn et al. (2021) also noted that soil aggregation, a critical aspect of soil health, is greatly influenced by microbial structures, functions, and plant interactions.

Furthermore, Azolla, known for its cost effectiveness and eco-friendliness, enriches soil with carbon and nitrogen (Akhtar *et al*., 2020). Additionally, a study by Ju et al., (2018) revealed that soil microorganisms such as Bacillus subtilis, Thiobacillus thioxidans, and Saccharomyces spp. can symbiotically fix atmospheric nitrogen, potentially satisfying 80–90% of the nitrogen requirement for crops through symbiosis with soybeans.

Biofertilizers containing soil microorganisms play a crucial role in enhancing soil fertility. In addition, their application enriches the structure of the soil and reduces the reliance on chemical fertilizers. For instance, research by (Imtiaz *et al*., 2016) and (Chaer *et al*., 2011) highlights the potential of bacterial and fungal inocula, along with organic amendments, as viable options for integration into crop-based nutrient management strategies aimed at improving degraded soils.

According to (Bardgett and Straalen, 2008), arbuscular mycorrhizal (AM) fungi and biological nitrogen-fixing bacteria collectively contribute 5-20% annually to the total nitrogen demand in grasslands and savannahs. In temperate and boreal forests, AM fungi alone contribute 80% to nitrogen acquisition, while bacteria and fungi contribute 75% to the acquisition of phosphorus by plants.

Soil flora and fauna play a crucial role in improving soil texture, nutrient levels, and crop productivity. Mycorrhizal fungi, for example, produce a sticky sugar protein called glomalin, which aids in the aggregation of soil particles due to its cementing properties (Marisângela Viana Barbosa, Curi, and Carneiro, 2019). Furthermore, Maanoja et al. (2020) found that bacterial decomposition of organic matter improves soil porosity, thus improving infiltration capacity and reducing the risk of erosion.

Arbuscular mycorrhizal fungi hyphae are highly effective in accessing inorganic phosphorus (P) within soil pores, thus improving the translocation of inorganic P and addressing soil P deficiencies (Bago, Pfeffer, and Shachar-Hill, 2000). Continuous use of blue-green algae biofertilizers has been shown to not only maintain but also increase the organic carbon content of the soil by up to 22% (Kumar et al., 2023). Additionally, co-inoculating effective rhizobia inoculants with biofertilizers aimed at improving P availability and uptake can enhance the efficiency of biological nitrogen fixation.

In China’s arid saline soils, deficiencies in phosphorus (P) and potassium (K) are common. However, the application of P-solubilizing bacteria has shown effectiveness in improving nutrient availability (Singh, Singh, and Chinna, 2023). Similarly, the integration of rock phosphate with P-solubilizing microorganisms has been found to improve the availability of P (Khan et al., 2009). In India, the effectiveness of chemical P fertilizers has been enhanced by using phosphate solubilizing bacteria (PSB), which has led some companies to advocate for increased sales of chemical fertilizers in conjunction with biofertilizers (Naseem, Nagarajan, and Pray, 2023).

Microbial polysaccharides play an important role in the formation and stabilization of soil aggregates (Costa, Raaijmakers, and Kuramae, 2018). Regular application of effective strains of blue-green algae (BGA) in soils can help decrease soluble salt levels, neutralize soil pH, and reduce sodium content in the exchange complex (Carillo et al., 2020). The combined effect of lowering soil pH, electrical conductivity, and exchangeable sodium, along with improving soil aggregation and permeability to air and water, increases soil carbon content, ultimately leading to improvements in soil health and productivity.

Bacteria and fungi play a crucial role in improving soil structure by aiding in the formation of soil aggregates and pores. Al-Maliki and Breesam (2020) observed that fungal cells produce mucilaginous exudates primarily made up of extracellular surface polysaccharides, cell wall polysaccharides, and intracellular polysaccharides. These extracellular polysaccharides are instrumental in the formation of aggregates, thus improving soil porosity and aeration. Additionally, bacteria produce exopolysaccharides that create organomineral complexes, which help bind soil particles to aggregates (Kaur et al., 2022).

Research shows that soil structure is influenced not only by mineral components but also by microorganisms present in soil pores (Vasilchenko et al., 2022). Furthermore, organic exudates from bacteria, fungi, decomposed cells, and plant and animal residues enhance soil organic matter, particularly in organically managed soils. This increase in organic matter content subsequently leads to improvements in soil structure, functionality, and overall quality.

Arbuscular mycorrhizal (AM) fungus hyphal networks play a crucial role in enhancing soil aggregation through both direct and indirect mechanisms. Studies by Ren et al. (2022) and Erktan et al. (2020) have shown that AM fungi mycelium directly contribute to soil aggregation by forming extensive hyphal networks that bind soil particles together. Furthermore, research by Charlotte et al. (2021) indicates that these networks also facilitate soil particle alignment along expanding hyphae, further promoting the formation of aggregates.

Wilkes (2021) reports that about 80% of the glomalin protein is located within the hyphal walls of arbuscular mycorrhizal (AM) fungi, helping to transport nutrients and water. Glomalin also protects fungal hyphae and lipid-rich spores from drought and microbial attacks, contributing to the formation of soil aggregates through decomposed hyphae and the fusion of glomalin protein with minerals and organic matter (Avila-Salem et al., 2020). The hydrophobic nature of this protein increases the hydrophobicity of soil aggregates, while its adhesive properties help initiate and stabilize soil aggregates (Al-Maliki and Breesam, 2020). Glomalin acts as an adhesive agent, binding soil particles together (Rillig, Wright, and Nic, 2001).

Saprotrophic fungi release extracellular exudates that facilitate the creation of water-stable soil aggregates (Zheng, Morris, and Rillig, 2014). Likewise, extracellular mucilages, such as basidiomycetes and Trichocomaceae, help to bind soil particles, improving aggregate stability (Daynes et al., 2013).

### 4.4 Role of microbial activities in crop productivity

Soil microbial activity significantly increases crop productivity, as reported in 42.1% of the articles analyzed (see Figure 4). For example, Abdel-Fattah et al. (2012) found that soil inoculation with algae-enhanced microorganisms led to crop yield increases between 16.1% and 32.8% in various Indian regions. Plants depend on soil microbes, such as bacteria and fungi, for essential nutrient access. These microorganisms have the metabolic ability to break down and mineralize organic nitrogen (N), phosphorus (P), and sulfur (S) (Bonkowski et al., 2009; Richardson et al., 2009). This process releases inorganic forms of N, P and S, such as ammonium, nitrate, phosphate and sulfate, which plants readily absorb, facilitated by microbial activity (Bardgett and Straalen, 2008). In natural settings, these microbial nutrient transformations are crucial for plant growth and can sometimes limit ecosystem productivity (Schimel and Bennett, 2004).

In lowland environments, the use of blue-green algae (BGA) in combination with Azospirillum has been shown to significantly increase grain yield (Choudhary et al., 2024). Similarly, Ju et al. (2018) found that biofertilizers containing Azotobacter, Rhizobium and Vesicular Arbuscular Mycorrhiza led to the highest increases in wheat straw and grain yields, especially when used with rock phosphate as a phosphate fertilizer. Furthermore, Osei Micheal Banahene (2020) demonstrated that the application of P fertilizers with bio-fertilizers increased soybean yields by approximately 47% compared to a negative control in low-P soils in sub-Saharan Africa.

Beneficial soil microorganisms, which function as biofertilizers or symbionts (Nina et al., 2014), support crop production by improving nutrient solubilization, thus improving nutrient availability and uptake (Tahir et al., 2022). They promote plant growth by enhancing root architecture development (Schillaci et al., 2021) and provide beneficial traits such as increased root hairs, nodules, and nitrate reductase activity (Lee et al., 2020). Additionally, soil microorganism biofertilizers contain plant hormones such as indole acetic acid (IAA), gibberellins (GA) and cytokinins (CK) (Hassan et al., 2022), which improve photosynthesis, improve stress tolerance (Chi et al., 2010), and increase resistance to pathogens (Thamer et al., 2011), ultimately leading to better crop yields.

Biocontrol, a modern disease management strategy, can greatly benefit agriculture, similar to biofertilizers. For example, (Ouledali et al., 2018) found that soil microorganisms can improve zinc (Zn) and copper (Cu) uptake while protecting against root diseases, thus promoting plant growth and enhancing crop productivity. Trichoderma, a biocontrol agent, has effectively controlled root rot in mung beans and increased yields (Ju et al., 2018). Additionally, biofertilizers containing bacterial nitrogen fixers, solubilizing bacteria of phosphorus (P) and potassium (K), and specific microbial strains have significantly improved the growth, yield, and quality of certain plants (Mohammadi and Sohrabi, 2012).

Garg et al. (2016) found that *Rhizobium trifolii* inoculated with *Trifolium alexandrium* showed higher biomass and more nodulation under salinity stress. *Pseudomonas aeruginosa* is known to endure biotic and abiotic stress. Ayub et al. (2024) reported that *Pseudomonas putida* improved the germination rate and various growth parameters, such as plant height, fresh weight and dry weight of cotton, in alkaline and high-salt conditions by increasing the uptake of K^+^, Mg^2+^, and Ca^2+^ while reducing Na^+^ absorption. Some Pseudomonas strains help plants tolerate soil stress (Mora et al., 2022). Mycobacterium phlei helps to tolerate high temperatures and salinity stress (Delgadoa et al., 2024). Tufail et al. (2018) demonstrated that arabuscular mycorrhiza fungi enhance plant growth under salt stress, and the root endophytic fungus Piriformospora indica helps protect host plants against salt stress.

Arbuscular mycorrhizal (AM) fungi combined with nitrogen-fixing bacteria help legume plants manage drought stress. Applying Pseudomonas sp. to plants during water stress increases their antioxidant levels and photosynthetic pigment content, improving seedling growth and seed germination (Sabeti et al., 2019). Furthermore, Aroca et al. (2010) observed that rice plants inoculated with AM fungi showed improved photosynthetic efficiency and antioxidative response under drought conditions.

Soil microorganisms not only promote growth, but also confer pathogen resistance through metabolite production (Kumar et al., 2015). Bacillus subtilis, for example, can activate defense pathways and when combined with mature compost, it effectively controls Fusarium infestation in banana roots (Yahuza and Tijjani, 2024). Patil et al. (2022) emphasized that rhizobacteria that promote plant growth (PGPR) help manage viral diseases in crops such as tomatoes, cucumbers, peppers, and bananas. Furthermore, mycorrhizae and bacteria together provide resistance against fungal pathogens, inhibiting the growth of root pathogens such as Rhizoctonia solani and Pythium sp (Lahlali et al., 2022).

Rhizobium is essential for increasing crop yields by converting atmospheric nitrogen into forms plants can use (Lee et al., 2020). Research by Arora and Sahoo (2015) demonstrated that Rhizobium inoculants significantly boost grain yields in Bengal gram in various locations and soil types. Additionally, Rhizobium strains isolated from wild rice have been shown to promote growth and development in rice plants by providing nitrogen (Padukkage et al., 2021). The Rhizobium species Sinorhizobium meliloti 1021 has been noted to infect nonleguminous plants such as rice, enhancing growth by increasing plant hormone levels and photosynthetic efficiency, which improves stress tolerance (Anli et al., 2020). In groundnuts, the IRC-6 strain of Rhizobium has been found to increase the number of

pink nodules, the activity of nitrate reductase and the content of leghaemoglobin 50 days after inoculation (DAI) (Lee et al., 2020).

Beneficial soil microorganisms that solubilize insoluble phosphorus forms are crucial to improving the efficiency of phosphorus use in agricultural soils. Research conducted in greenhouse and field environments (Ashraf et al., 2021; Gulshan et al., 2022; Ramrez-Flores et al., 2020) has shown that the use of phosphate-solubilizing microorganisms (PSM) and arbuscular mycorrhizal fungi (AMF) leads to improved phosphorus uptake and increased yields in various vegetable and cereal crops.

Research has shown that inoculating various legumes, such as cowpea, chickpea, soybean, and common bean, with rhizobia and supplementing with phosphorus fertilizer improves the uptake of essential nutrients such as nitrogen, phosphorus, potassium, magnesium, calcium, and sodium (Rotaru, 2018; Al-Burki & Al-Ajeel, 2021). Inthasan et al. (2023) observed that mung bean inoculated with Rhizobium and provided with phosphorus (45 kg P2O5 ha-1) showed improved symbiotic performance, including increased biomass, number of nodules, and nitrogenase activity. Furthermore, Kyei-Boahen et al. (2017) found that combining rhizobia inoculants with mineral phosphorus fertilizer improved the agronomic efficiency of cowpea-Bradyrhizobium symbiosis compared to using either treatment alone. The application of phosphorus also enhances the effectiveness of indigenous rhizobia, leading to a higher nitrogen content in shoots and seeds, as well as to improved grain yield and plant biomass.

The combination of phosphate-solubilizing microorganisms (PSM) with phosphorus has been extensively used, showing significant improvements in the agronomic efficiency of rock phosphate and various phosphorus fertilizers such as DAP, NPK, and TSP (Duarah et al., 2011; Kaur and Reddy, 2015). Duarah et al. (2011) demonstrated that the use of NPK fertilizer along with a consortium of seven PSB strains, known for their high phosphorus solubilization abilities (including Staphylococcus epidermidis, Pseudomonas aeruginosa, Bacillus subtilis, and Erwinia tasmaniensis), resulted in improved plant biomass and increased germination indices in rice and cowpea beans. These benefits were linked to the stimulation of enzyme biosynthesis, such as amylase, in seeds.

Additionally, a study by Noor et al. (2017) examined maize growth when DAP fertilizer was impregnated with Pseudomonas putida. The DAP was coated (20 g/kg) with a mixture of organic materials, including compost, molasses, and the strain of *P. putida*. The results indicated that the combination of DAP with PSB improved the yield of dry matter from maize by 12% and the uptake of phosphorus by 33%, significantly improving the agronomic efficiency. Biomass production saw a 62% increase compared to unfertilized soil. This method of integrating P fertilizer with bacteria offers a promising solution for challenging soil phosphorus management. Further research on efficient rhizobial isolates and strains in various Ethiopian agroecologies suggests that biological nitrogen fixation (BNF) can substantially increase food production by increasing cowpea crop and forage yields. For example, Hailemariam & Angaw Tsigie (2003) reported crop yield increases between 51% and 158% on Nitosols in Holleta, Ethiopia, when 20 kg P ha-1 was applied with the strain, compared to noninoculated crops.

## 5. Conclusion and Implications of the Policy

The soil is home to a diverse range of organisms that interact with plants in a complex way, primarily through nutrient cycling and transfer. Various studies highlight the significant role of soil microorganisms in this process, as they are crucial in recycling both macro and micronutrients. Furthermore, soil microbial activities are essential in converting organic and inorganic nutrients into forms that crops and microorganisms can easily absorb.

Biological components in the soil are essential to improve and maintain soil fertility. The application of biofertilizers significantly enhanced the physical and chemical properties of the soil. An increase in nutrient availability was observed after soil inoculation with fungi or bacteria. Numerous studies emphasize the critical role of soil microorganisms in crop production in various regions. Inoculation of the soil with these microorganisms results in substantial increases in yield and yield components for a variety of crops.

The review highlights the importance for farmers to manage soil microorganisms to optimize soil and crop production. Sustainable agricultural practices depend on integrated management of land resources by policymakers, experts and managers, with a focus on improving soil microorganism health. The current review primarily examined specific soil microorganisms, such as bacteria and fungi. Future reviews should expand their scope to include other components of the soil fauna and flora.

## Conflict of Interest

The authors declare that they have no conflict of interest.

## Notes

### Competing Interest Statement

The authors have declared no competing interest.

